# Optimal compositions of potent sliced-siRNA with DNA bases substitution and insights of Ago2 slicing actions

**DOI:** 10.64898/2026.11.03.764113

**Authors:** Guihua Sun, Elizabeth M. Lizhar, Yanhong Shi, Arthur D. Riggs

## Abstract

We have previously characterized the nucleotide compositions of single-stranded ago2-sliced small interfering RNAs (sliced-siRNA, sliRNA), a novel type of potent RNAi trigger designed based on the biogenesis mechanism of miRNA-451. sliRNAs have on par antisense mediated silencing activity but about 1000-fold reduced passenger strand activity compared to siRNA. In the current study, to further characterize sliRNAs to facilitate their *in vivo* application, we have investigated the effects of substituting RNA nucleotides (nts) in sliRNA with DNA nts (sliDRNAs), which will allow us to study the potential effects on their processing, maturation, and silencing activity from individual or multiple nts at different positions. Our results showed sliDRNAs have distinguished composition from siRNA with DNA nts (siDRNA), indicated that, in addition to their different biogenesis mechanisms, these two types of RNAi triggers may also have adopted additional different silencing mechanisms. Our results also revealed some mechanism related to how Ago2 executes its slicer activity and the effects on sliRNA potency by trimming and tailing during sliRNA maturation. These results have paved the way for using other chemically modified nts in sliRNAs that may enhance their *in vivo* stability and transportability to facilitate their *in vivo* applications.

## Introduction

RNA interference (RNAi)—using small interfering RNAs (siRNAs), microRNAs (miRNAs), or antisense oligonucleotides (ASOs) to silence gene expression—is a powerful gene manipulating strategy^1–6^. Their ability to target virtually any gene in a pathway by a natural endogenous mechanism makes them ideal reagents with great potential for treating human diseases^7,8^. However, the application of RNAi reagents, particularly *in vivo*, faces several major challenges, including instability due to RNases-mediated degradation, rapid renal clearance due to their small mass size, and inability to penetrate cell membrane on their own without effective delivery reagents. Other potential problems are off-target effects (off-TEs), innate immune stimulation, guide/passenger strand (GS/PS) separation issue during delivery, and limitation of choosing a potent GS^3,4^. Fortunately, the miR-451 biogenesis pathway has provided us with a template from which we have designed a novel type of RNAi trigger, ago2 sliced-siRNA (sliRNA), may overcome some of the above challenges faced by traditional siRNA (**Fig. 1**) ^9–14^.

**Fig. 1.**
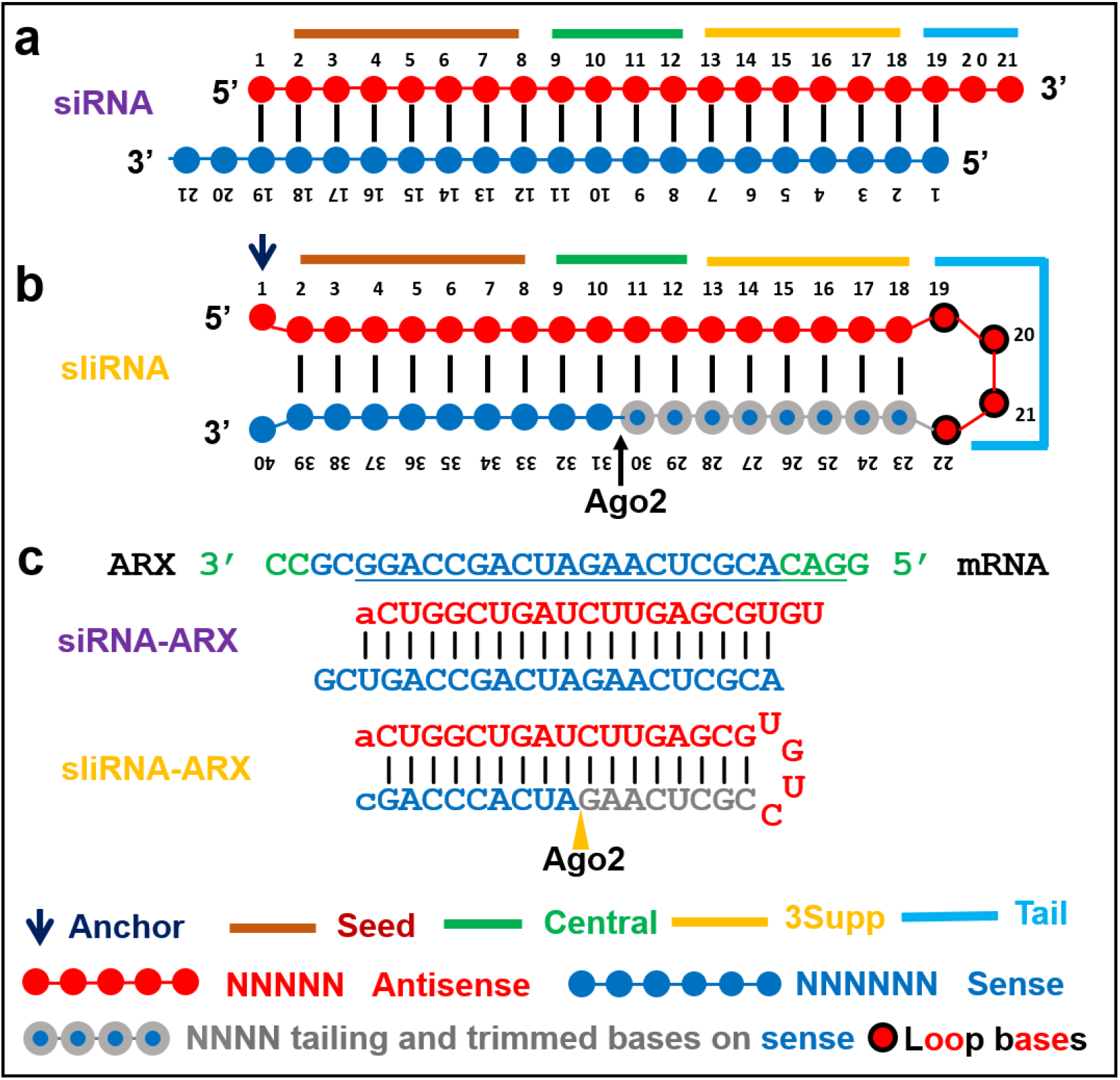
Schematic illustrations of **a)** a siRNA, consisting of a 21-nt passenger strand (sense) and a 21-nt guide strand (antisense); **b)** a sliRNA, consisting of an anchor nt, a 17-nt stem, the last 4 nts of the antisense strand as a loop, and the unpaired last nt (N_40_); **c)** An example of a siRNA and its corresponding sliRNA target the same region on the *ARX* gene mRNA. The 1^st^ nt on guide strand was changed from ‘C’ to ‘A’ to facilitated guide strand loading onto Agos.

Classical siRNAs, mainly defined as an asymmetric duplex RNA of 21 nucleotides (nts) with 2-nts 3’ overhang. The strand that performs RNAi is defined as antisense (AS) or GS; the other strand usually called sense strand (SS) or PS^15^. siRNAs will be incorporated into Argonautes (Agos, Ago1 to Ago4 in mammals) to form RNA-induced silencing complex (RISC) to perform RNAi function^7,16,17^. If the AS is fully complementary to the target RNA, and they are in Ago2-loaded RISCs (Ago3 loaded RISCs under some special conditions), RISCs will cleave the target RNA for target degradation, if they are in non-slicing Ago-loaded RISCs (Ago1, 3, and 4), RISCs will block target RNA translation or trigger target RNA degradation. If the AS is partially complementary to the target RNA, all RISCs will mainly block target RNA translation or trigger target RNA degradation. Because RISC preferentially select the strand with the less stable 5’ end to form the active complex, siRNA need to be designed as duplex with differential ends thermal dynamic energy, lower thermal dynamic stability at the 5’ end of the GS and greater stability at the 5’ end of the PS^18^, to favor GS loading over PS and to achieve the intended on target effects (on-TEs). Otherwise, more RISCs will be loaded with the PS that can cause unwanted, maybe toxic, off-TEs^19,20^. This requirement thus restricts sequences that can be used as GS for siRNA. One solution to reduce PS-mediated off-TEs is to use oligonucleotides of single-stranded AS alone without the SS. But RISCs have more than thousand-fold higher binding affinity for duplex RNAs over single-stranded RNAs^21^.

Traditional siRNAs are mimics of DROHSA then Dicer produced miRNA/miRNA* mimics. Nature also evolved out a Dicer bypassing mechanism to generate mature miR-451^9–11,22–24^. Drosha-produced precursor miRNA-451 (pre-miR-451) is a small hairpin structure of the 5p strand (N_01_-_18_) physically linked with the 3p strand (N_23_-_42_) by a small tetra nts loop (4-nt loop, N_19_-_22_). The hairpin structure has a short stem region (N_02_-_18_:N_23_-_39_) and a small loop that it cannot be processed by Dicer, instead, Ago2 will cut in the middle of the 3p (between N_30_ and N_31_) to generate a longer fragment of N_01_-_30_ and shorter resected fragment of N_31_-_42_, then the N_01_-_30_ fragment will be trimmed by Poly(A)-Specific Ribonuclease (PARN) from 3’ end to form the 22-nts mature strand (N_01_-_22_)^25,26^(**Fig. 1b**). We have extensively characterized pre-miR-451 mimics and formed the optimal sliRNA nts composition^14^.

sliRNA has target cleavage and repression activity comparable to that of classical siRNA and several potential advantages over siRNA^14^. First, sliRNA has 100-to 1000-fold less PS activities that will reduce off-TEs due to its inherited property from pre-miR-451 that the GS maturation must destroy the PS. Secondary, theoretically sliRNA can be implemented with any sequences as GS due to 5p of the hairpin is predefined as GS. In siRNA, the GS is selected in RISC according to the differential ends thermodynamic properties of the GS/PS duplex, which limits antisense strand selection for a specific gene. This limitation is more severe for genes with short sequence, or for genes that are specific isoforms. Third, being a single molecule, sliRNA may avoid the strand separation risk associated with siRNA delivery and may also facilitate their delivery by attaching delivery groups to the last nt at 3’ end. Despite these advantages, sliRNA have not been used as an *in vivo* RNAi trigger, mostly because of lacking studies of their *in vivo* stability and deliverable ability.

*In vivo* used siRNAs need to be heavily modified by varies chemical modifications on nts, including modifications on phosphate backbones, ribose rings, and bases. These chemical modifications are mainly used to improve siRNA nuclease resistance for higher stability. It can also be used to increase target RNA binding affinity (higher melting temperature) for effective binding and long-lasting silencing. Another important benefit of chemical modification is that it can modify charges of RNAi reagents to increase overall hydrophobicity of RNAi molecules, which are negative charged using natural RNA bases, to enhance their ability to penetrate cell membrane or package into delivery reagents. Last, chemical modifications were used to reduce immune stimulation by specific RNA motifs within a siRNA molecule^27–31^. Common modifications on phosphodiester (P=E) backbone are phosphorothioate (P=S) and phosphotriester (P=TE). Most modifications made on nucleobases are on uridine, such as 5’-Bromo-, 4-Thio-, N3-Methyl-, 5’-Iodo-uridine, etc. Most widely used and effective chemical modifications to siRNA are done on the sugar ring of nucleosides, such as 2’-Deoxy (DNA), 2ʹ-O-methyl (2’-OME), 2’-Deoxy-2ʹ-Fluoro (2ʹ-F), 2ʹ-O-methoxyethyl (2ʹ-MOE), locked nucleic acid (LNA), etc^8^. In addition, specific chemical moieties are attached to GS or PS to facilitate packaging or delivery. In theory, the effect of chemically modified nts (cm-nts) on sliRNA and siRNA will be different due to their RISCs use different functional mechanism for silencing function^25,32^. There is a potential problem with cm-nts on PS of a cm-sliRNA. Activation of a siRNA with cm-nts (cm-siRNA) loaded-RISC (siRISC) can use PS cleavage or ejection mechanisms and cm-nts on PS may help reduce PS activity ^18,33,34^. But a sliRNA with cm-nts (cm-sliRNA) loaded-RISC (sliRISC) needs Ago2 to slice the 3’ arm (equal to the PS in a siRNA) to be activated and Ago2 may not be able to cut PS with cm-nts. Therefore, cm-nts probably will have negative effect on potency of cm-sliRNA. To answer these questions and to facilitate sliRNA *in vivo* application, we have performed systematic DNA nts substitution on sliRNA to determine the optimal DNA nt replacements for which the following are minimally affected: silencing function (Ago2 processing, sliRISC formation and activation), off-target effects, and immune activation, as an economic way to forecast the composition of cm-sliRNA.

Replacing RNA nts with DNA nts in siRNA is an attractive approach to test cm-siRNA due to DNA nts are more stable than RNA nts, cost effective (10 times cheaper), and most importantly, DNA nts are also negatively charged and mainly different from RNA nts at the 2’ group on the sugar ring^33,34^. Other benefits of DNA replacements in siRNA are reduced seed-mediated off-TEs, reduced PS activities, and reduced immune stimulation effects. Therefore, DNA nts replacement in siRNA is an easy and convenient way to study the potential effects on siRNA activity from chemical modifications on nucleoside. In the current study, our results showed compositions of potent sliRNA with DNA nts (sliDRNA) are different from siRNA with DNA nts (siDRNA), implied that slicing-competent Ago2 may exhibit different conformations in sliRISC versus siRISC, and trimming and tailing on processed sliRNAs may affect their potency. Furthermore, more nts can be replaced in sliDRNA than siDRNA to further reduce production costs and extra nts can be tailed at sliRNA end to make multiple function chimera.

## Results

### Different composition of sliDRNA versus siDRNA

Previous studies in siDRNA showed the anchor base (nts #1 on GS, g1), base-paired nts in the seed region (g2-g8 on GS and p12-p18 on PS), p19-p21 on PS, and g13-g14:p6-p7 pairs in the 3’ supplementary region (3supp) in a siRNA duplex can be substituted with DNA nts to reduce seed-mediated off-TEs (**Fig. 1a**, **2a**), immune stimulatory effects from certain RNA motifs, and production cost while maintain its potency^33–35^. To validate the above findings and to test whether the same DNA replacements strategy used on siDRNA can be adapted to sliDRNA, we have compared DNA nts substitutions in siRNA-ARX and sliRNA-ARX (target human *ARX* gene). (**Fig.1c**, **2a**). We found **a**) Both siDRNA and sliDRNA reagents can be highly potent. In general, our data agreed with published results that siDRNAs and siRNAs have similar target knockdown rate at high dosages^33,34^. However, at lower dosage, DNA base substitutions will reduce potency for both types of RNAi reagents, and si/sliDRNAs were less potent than si/sliRNAs at lower concentrations. The same DNA substitutions will be more negatively affect sliRNA potency than siRNA (**Fig. 2b**: si-versus si-DR-I and -II; sli-versus sli-DR-I and -II); **b**) Although both siRNA and sliRNA can tolerate DNA bases in the seed region (DR-I) and retained well over 80% of parental molecules activities, combination of DNA base substitutions in both the seed region and 3supp (DR-II) reduce potency more than replacements in the seed region alone, implied, beside seed region, 3supp region are also critical for silencing function of RNAi reagents; **c**) DNA substitutions in 3supp region of sliRNA have much more effects than the same substitution on siRNA (**Fig. 2b**: si-/sli-DR-I vs -DR-II, si-DR-II vs sli-DR-II), implied siDRNA composition can’t be directly transformed into sliDRNA composition; **d**) Reduce DNA pairs in seed region has slight effects on sliDRNA (sli-DR-I vs -III and -IV), however, DNA substitutions on the anchor nt (N_1_) of GS have significant effects on sliDRNA potency (**Fig. 2b**: sli-DR-III/IV vs sli-DR-III/IV-R1), indicated that the anchor base play a critical role in sliRNA as well as in sliDRNA.

**Fig. 2.**
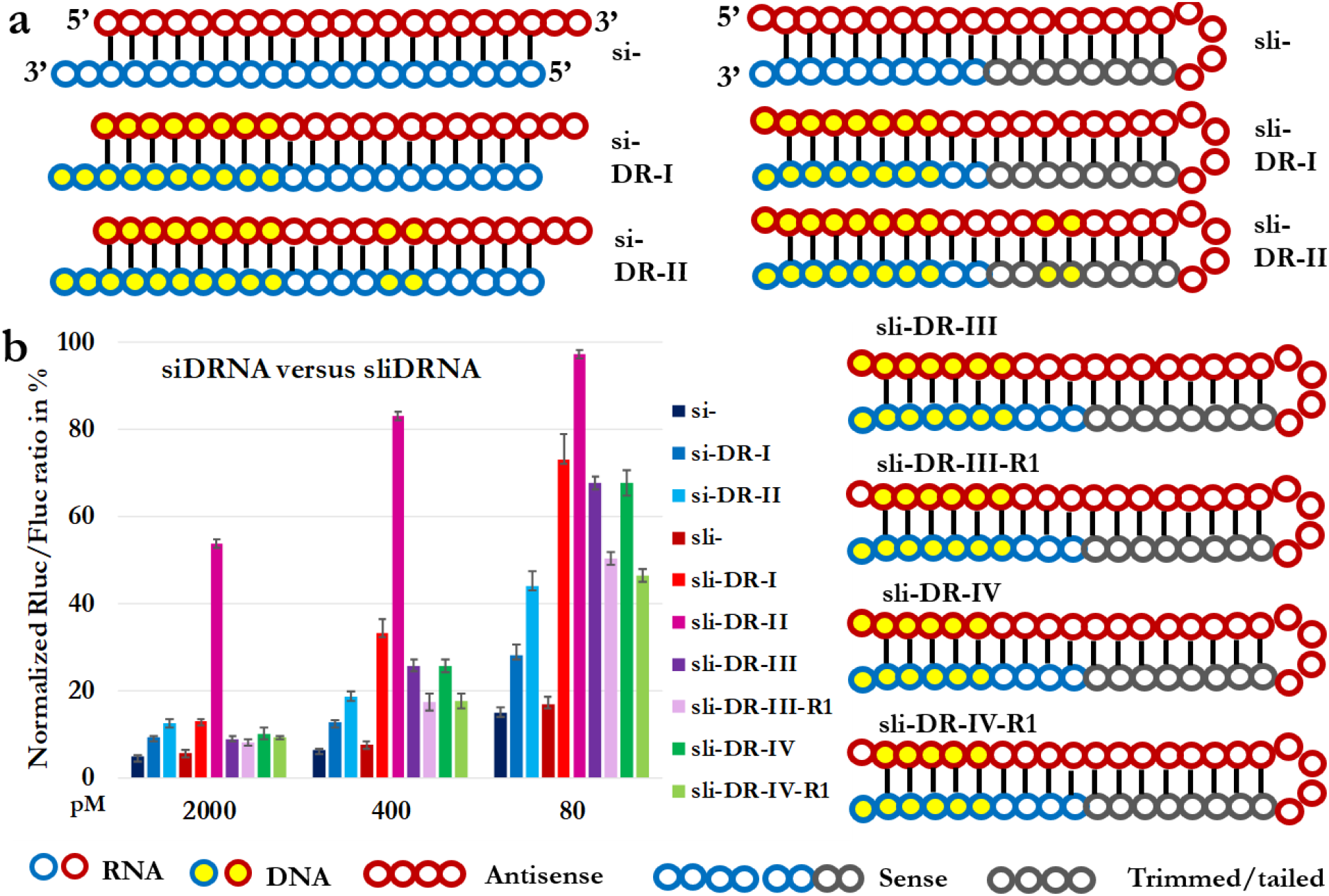
Compare potency of siDRNAs and sliDRNA designed based on the composition of siDRNAs (all RNAi reagents are based on a siRNA antisense sequence target human *ARX* gene): si-: siRNA; si-DR-I: composition #1 of siDRNA in reference; si-DR-II: composition #2 of siDRNA in reference; sli-: sliRNA; sli-DR-I: si-DR-I converted sliDRNA; sli-DR-I: si-DR-II converted sliDRNA; sli-DR-III and -IV are newly designed sliDRNAs and sli-DR-III-R1 and -IV-R1 are their corresponding sliDRNAs with N_1_ is a RNA nt. **a**. Schematic representation of all RNAi reagents used; **b**. Results from reporter assay.

The above results suggested, like siDRNA, sliDRNA can be potent, but it is unlikely the same DNA substitution rules used in siDRNA can be fully applied to sliDRNA. Systematical DNA substitutions on sliRNAs are needed to find the optimal potent composition for sliDRNA. To this end, we have performed systematical DNA substitutions on a sliRNA-ARX, then applied the rules to other sliRNAs to validate our finding (**Fig. 1c**).

We choose this sliRNA-ARX due to the two following reasons: First, we have extensively characterized this sliRNA before and it is a relatively less potent sliRNA compared to other sliRNAs target ARX^14,32^, therefore it will be easy to observe the positive or negative effects from DNA substitution on its potency; Second, there is undetectable ARX expression in HEK-293 cells using quantitated PCR, therefore HEK-293 cells is an adequate cell line for reporter assay that will have much less confounding effects from endogenous genes. Unless specified, all sliDRNA results generated in the following experiments, are from different forms of sliDRNA-ARX. We performed both individual and multiple DNA bases substitutions on sliRNA-ARX to find positions and regions that are critical for sliDRNA potency.

After Ago2 slicing the bond between N_30_ and N_31_ (on PS) to generate the large 30-nts fragment N_1-30_ and the smaller 10-nts resected (N_31-40_) fragment (**Fig. 1b**), the N_1-30_ fragment will be trimmed or tailed from N_30_ to generate many processed sliRNAs that are derivatives of the same sliRNA^9–11,14^. According to their alignment with parental sliRNA, we will name these processed isoforms as L_n_, with n representing the last nt that matches the n^th^ nts on the parental sliRNA (e.g., sequences match N_1-16,_ N_1-22,_ and N_1-30_ will be designated as isoform L_16_, L_22_, and L_30)_. Sequence length shorter than 15 nts will not be studied due to isoforms shorter than L16 have significantly reduced silencing potency^14^. Previously, we and others have proposed that it is possible that trimming/tailing actions maybe be happened while sliRISC perform silencing function using partial sequences from the processed partial hairpins (mainly L_23_ to L_26_) to base pair with targets^14,25,32^.

### DNA substitutions at the central region

At the central region, N_11_ is the nt used by Ago2 to slice the P=E bond between N_30_ and N_31_ to generate the large L_30_ and the small 10-nts resected fragment N_31-40_ (**Fig. 1b**). We found all single nt replacements from N_10_ to N_12_ can be tolerated and there is no significant effect. There is a slight effect from the replacements of all 3-nts from N_10_ to N_12_ (D_10-12_) or all 4-nts from N_9_ to N_12_ (D_9-12_). There is obviously an increased effect for replacements of all 6-nts from N_8_ to N_13_ (D_8_-_13_), all 8-nts from N_7_ to N_14_ (D_7-14_), and all 10-nts from N_6_ to N_15_ (D_6-15_), but D6-15 still remained over 70% activity. These results showed that Ago2 don’t need an RNA nt to execute the slicer activity and almost full slicer activities or over 80% activities will be retained after replacing 5 to 6-nts or 8 to 9-nts around N_11_ with DNA bases respectively. Despite this, when we replace most nts with DNA and only retain RNA at N_27-34_ (R_27-34_) or N_28-33_ (R_28-33_), both forms are completely inactive, indicated that RNA bases are important for sliRNA function. These results imply that, although Ago2 does not need an RNA to perform slicing function, RNA nts formed structure or environments will affect the slicer activities (**Fig. 3**).

**Fig. 3.**
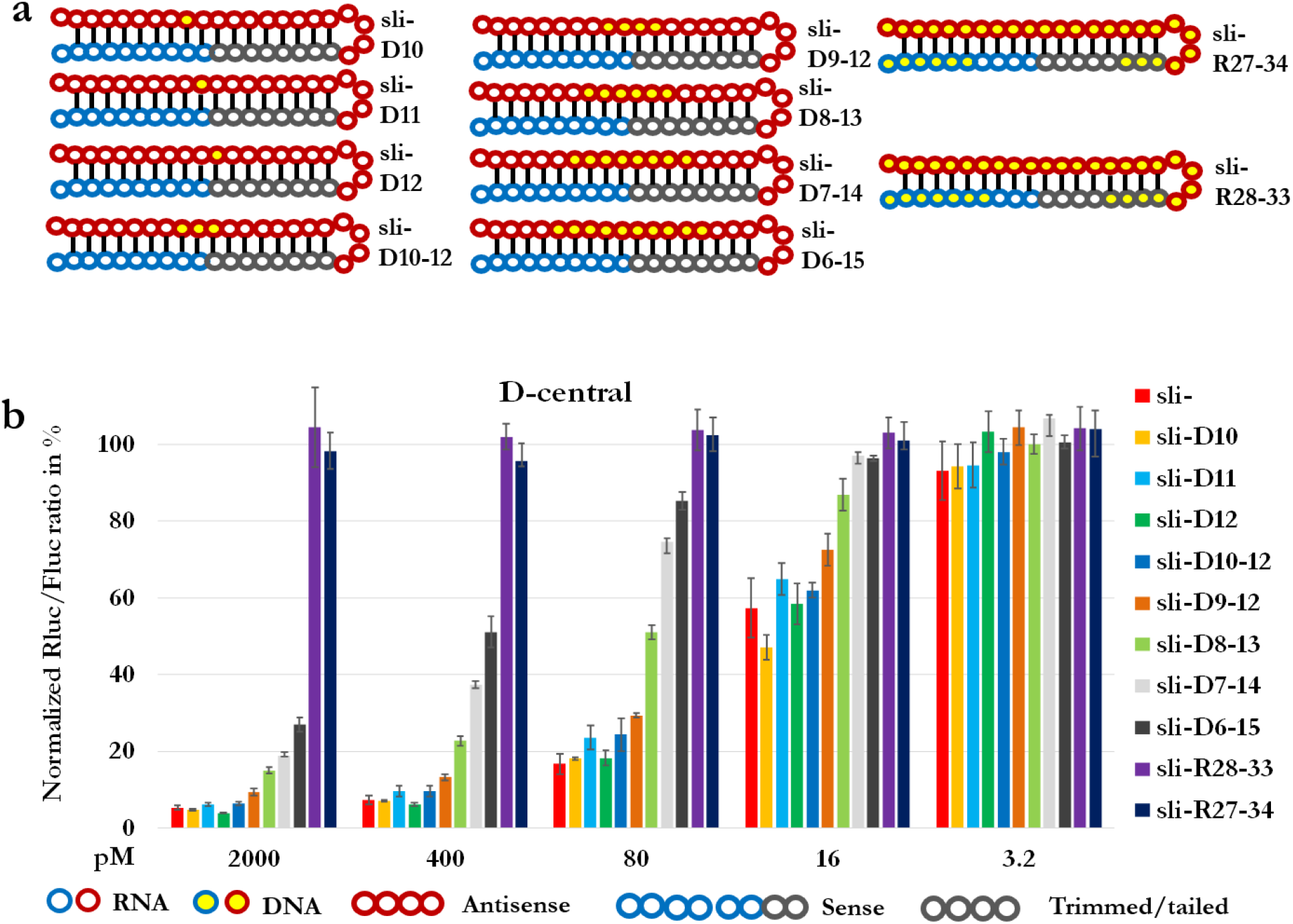
DNA replacements at central region. a. Schematic representation of all sliDRNAs; b. Results from reporter assay.

### DNA bases replacement at the trimming/tailing region (N_23-30_) and tetra loop (N_19-22_)

We performed DNA substitutions on the tetra nts loop region and found 1-nt (D_19_, D_20_, D_21_, and D_22_), 2-nts (D_21-22_), 3-nts (D_20-22_) or all 4-nts (D_19-22_) replacements do not have significant effects (**Fig. 4**). This agrees with our previous results that the small 4-nts loop can be swapped with other sequence without significant loss of silencing efficacy^14^. Next, we performed single DNA base substitution from N_23_ to N_29_ and found none of them has significant effects even though this region is used for trimming to produce mature sliRNA.

**Fig. 4.**
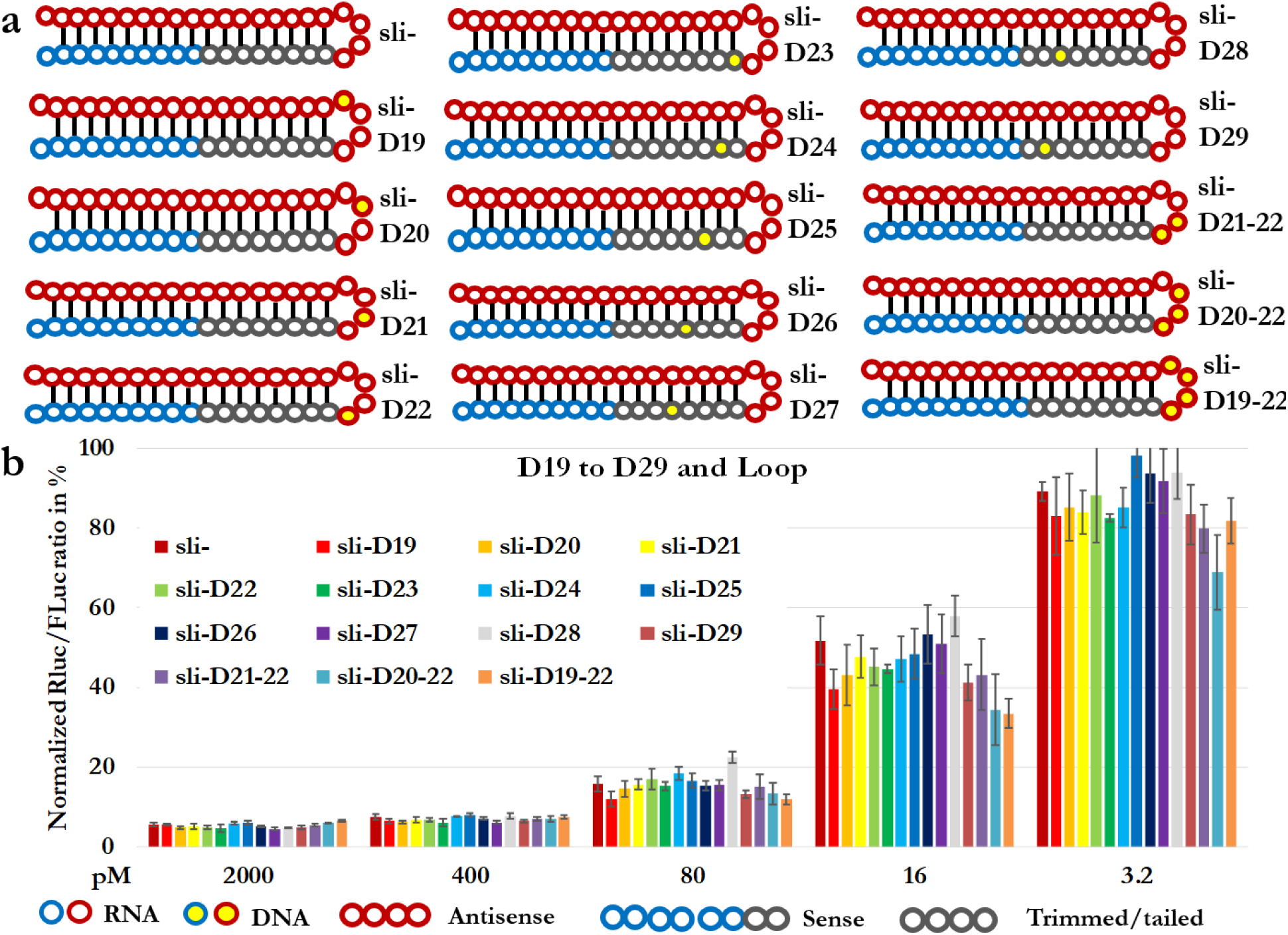
DNA replacements at N_19-29_ and loop regions. a. Schematic representation of all sliDRNAs; b. Results from reporter assay.

### DNA substitutions on the 3supp-loop region

Since the DNA replacements at the 3supp region have stronger effects on sliDRNA (**Fig. 2** sli-DR-II) and single nts replacements at N_19-29_ region have no significant effects (**Fig. 4**), we performed multiple base pair replacements at N_13-28_ region. We found, among all three regional replacements, 4-pairs (D_15-18:23-26_), 5-pairs (D_14-18:23-27_), and 6-pairs (D_13-18:23-28_), only the 4-pairs replacements, without the base pairs of N_13:28_ and N_14:27_, still retain over 80% potency (**Fig. 5**). These data support the previous observation that N_13:28_ and N_14:27_ pairs are critical for sliDRNA potency (**Fig. 2** sli-DR-II). In the combination of replacements at both 3supp and loop regions, we also found the base pairs of N_13:28_ (D_13-28_) and N_14:27_ (D_14-27_) replacements have more effects whereas loop region do not have significant effects, and one replacement at both 3supp and loop regions, the D_15-26_ form, still can maintain high potency (**Fig. 5**).

**Fig. 5.**
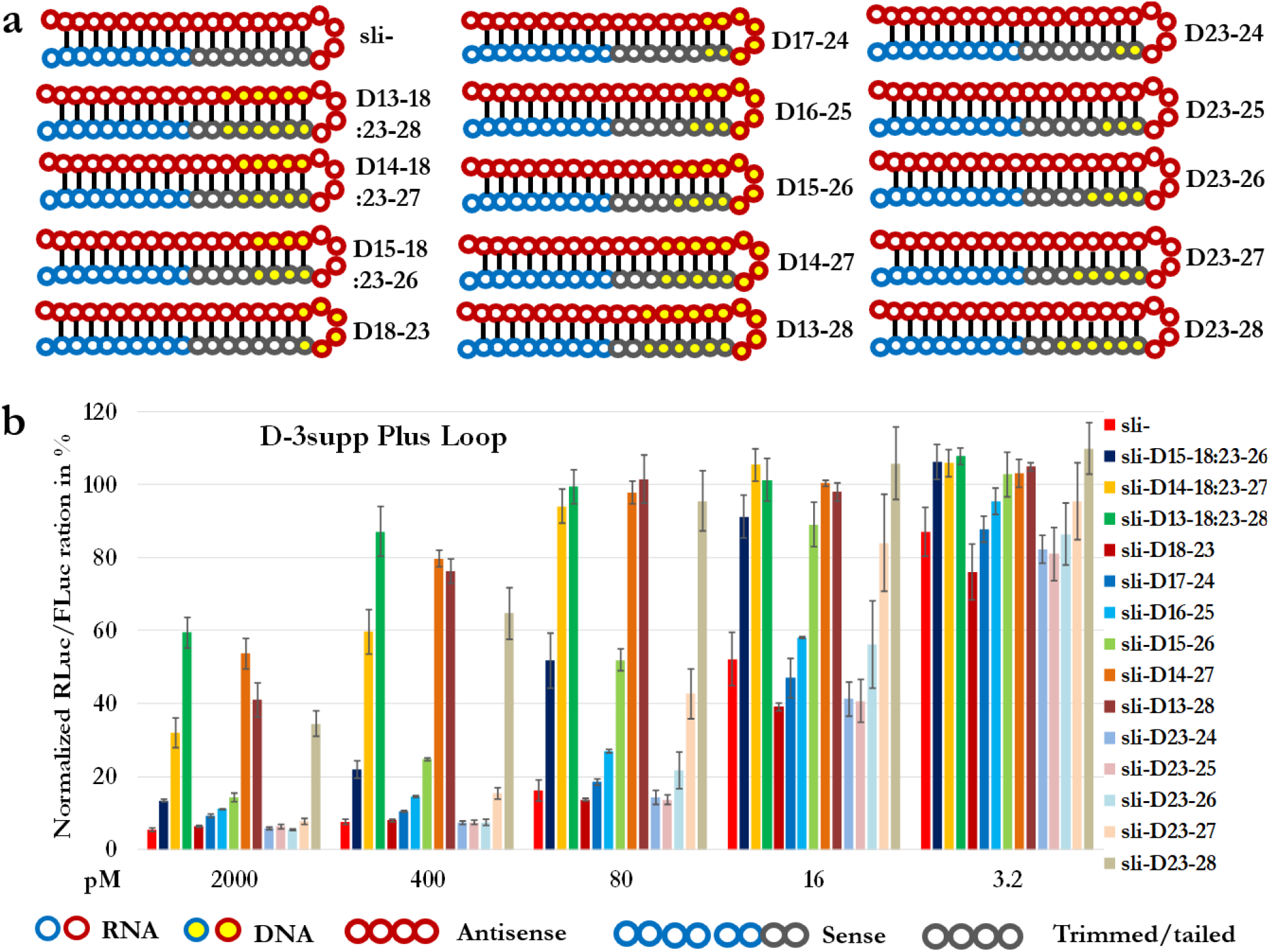
DNA replacements at 3supp-loop and the trimming-tailing nts. a. Schematic representation of all sliDRNAs; b. Results from reporter assay.

### Multiple DNA substitutions on the trimming-tailing nts

We found, the 4-nts-replacement at N_23-26_ does not have significant effect, however, the 5-nts-replacement (D_23-27_), which including N_27,_ starts to show reduced potency at lower concentration, and the 6-nts-replacement (D_23-28_), which including both N_27 and_ N_28_, greatly reduce sliDRNA potency to almost nonfunctional level (**Fig. 5**). This data further proved substitutions that include N_27_ or N_28_ will greatly affect sliDRNA potency and base-pairings at N_13:28_ and N_14:27_ are critical for sliRNA potency.

### DNA replacements at the N_30_ to N_40_ region

For sliRNA, it is necessary for 5p to slice 3p to activate sliRISC and 3p also acts as a target. However, single DNA replacement at N_30_ (D_30_) or N_31_ (D_31_) plus replacement of the 4-nts loop showed no effects at all (**Fig. 6**), indicated both the slicing bases (N_10_ and N_11_) and sliced bases (N_30_ and N_31_) do not need to be RNA bases. We further tested bases around N_30_ and all Ago2 resected bases (N_31-40_). We found all the bases from N_29_ to N_40_ can be replaced with DNA bases (Replacements of N_40_ with other bases were performed in our sliRNA composition paper^14^) with few percentage drops of potency at high concentration (2 µM). However, here again, any replacements involved N_27_ and N_28_ (base pairing with N_14_ and N_13_ respectively) will have much larger effects (D_27-32, -34_, D_28-32, -33_), but replacements at N_29_ and beyond have much smaller effects. Based on these data we can form a R25D15 form of potent sliDRNA (R_1_-_18_D_19-22_R_23-28_D_29-40_) that can reduce the cost of sliRNA by about 40%. Since we only observed minor effects from DNA nts replaced loop, we started to use sliRNAs with 4-DNA-nts loop for subsequent experiments.

**Fig. 6.**
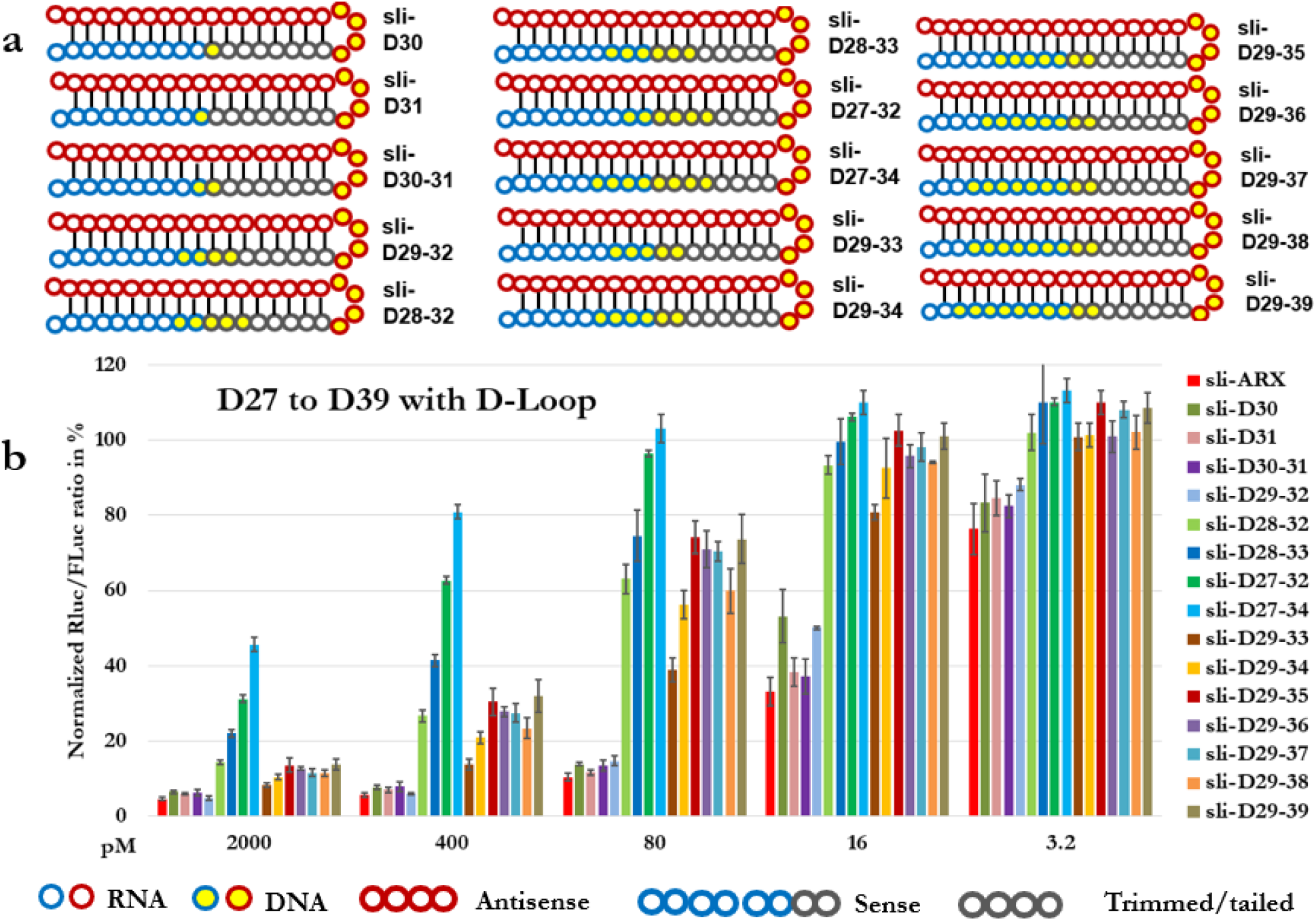
: DNA replacements at N_27-39_ with D-Loop. a. Schematic representation of all sliDRNAs; b. Results from reporter assay.

### Paired DNA replacements at the central region

Next, we tested paired DNA replacements around N_11_. Our data showed they can tolerate three DNA base pairs N_10-12:29-31_. Even though it can tolerate D_9-12_ or D_29-32_ alone, 4-nts DNA replacements around slicing bases or sliced bases respectively, the combination of 4 paired replacements started to show larger effects (**Fig. 7**).

**Fig. 7.**
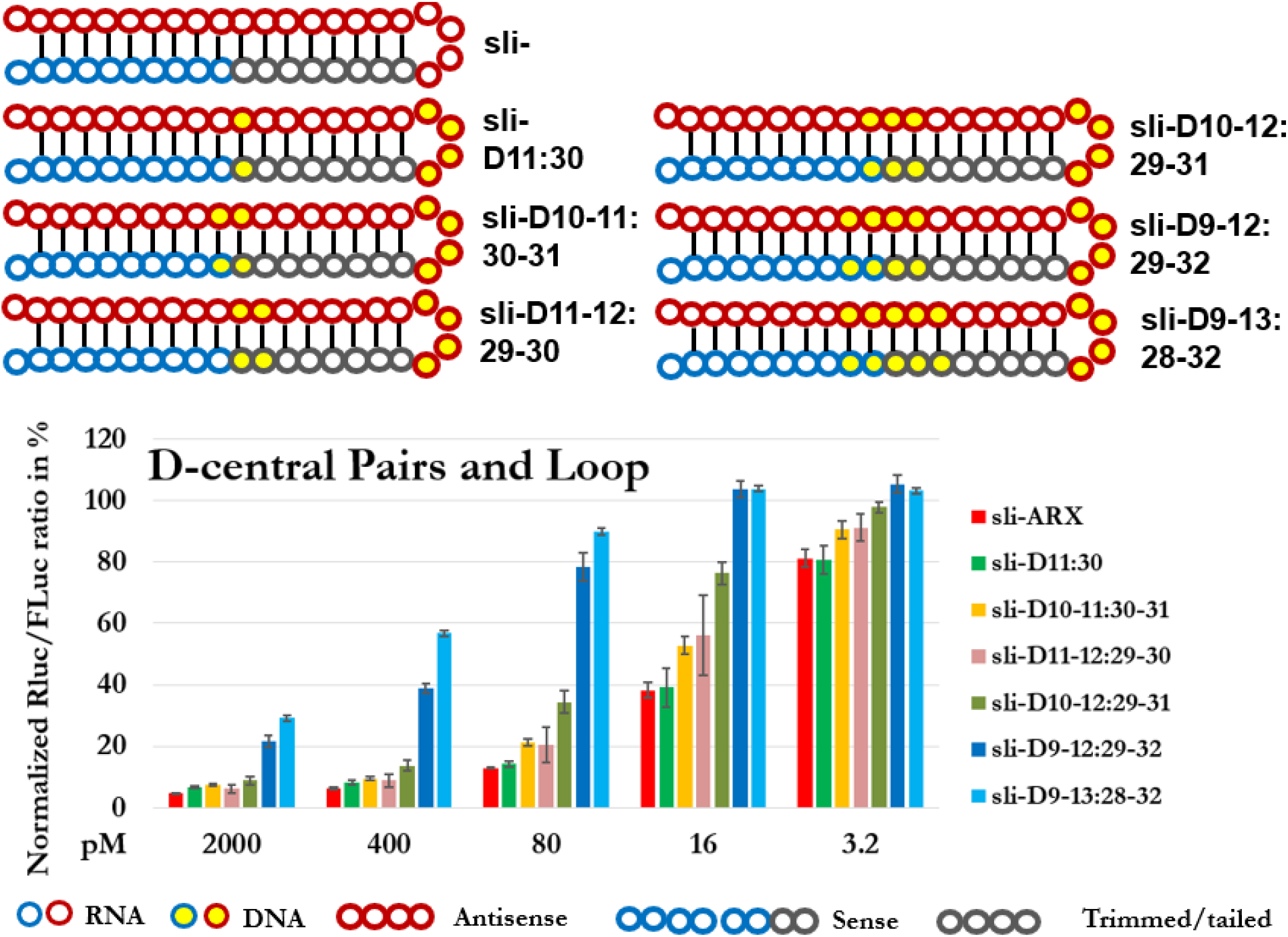
: DNA replacements at central pairs with D-Loop. a. Schematic representation of all sliDRNAs; b. Results from reporter assay.

### sliDRNA composition of DNA replacements at seed-central-loop regions

Based on all above experiments results, we started to form a potential sliDRNA composition with DNA replacements at seed, central, and loop regions (Dscl) that we can optimize later. We made three versions of Dscl (Dscl, Dscl2, and Dscl3) and tested using DNA or RNA as anchor base for version 1 and 2 (Dscl-R_1_, Dscl2-R_1_). We found only sli-Dscl-R1 can retain about 80% of sliRNA activities and further proved the critical role of RNA anchor base for sliDRNA (**Fig. 8**).

**Fig. 8.**
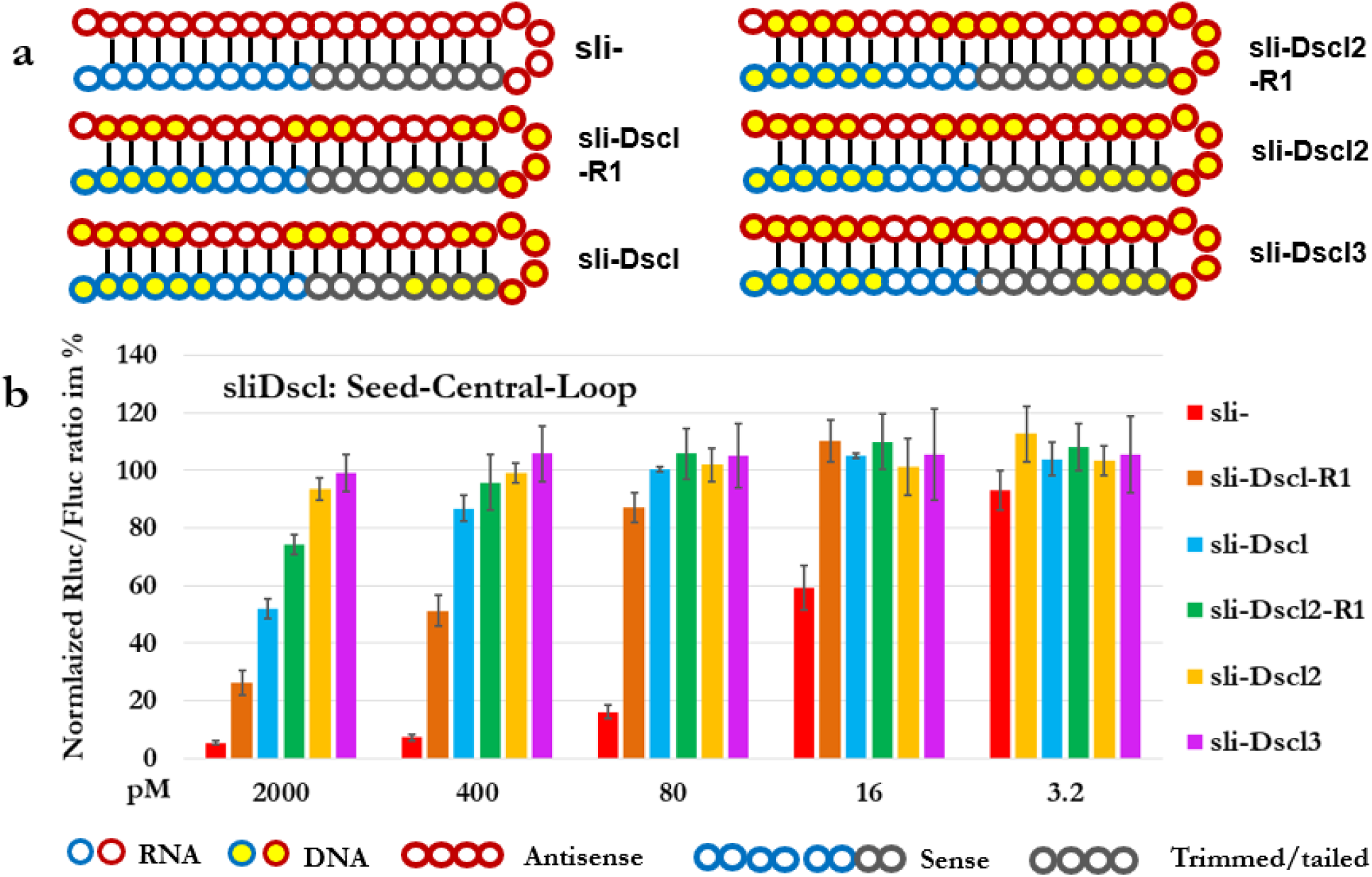
: sliDRNA composition of DNA replacements at seed-central-loop regions. a. Schematic representation of all sliDRNAs; b. Results from reporter assay.

### Candidate compositions of sliDRNA

Summarizing and combining all the above DNA replacements results, we formed three provisional compositions of sliDRNA, denoted as C1, C2, and C3 (**Fig. 9**). We tested their potency for sliDRNA-ARX, -STAT3 (−1623), -TLX (−934), -RRM2 (−1354), and -451 (sliRNA of miRNA-451) by reporter assays. Our results showed for all five sliDRNAs, both C1 and C2 performed significantly better than C3, but all three forms showed lower silencing activities than their corresponding sliRNA form, the difference is more obviously at lower concentration. However, the highest concertation we have used in reporter assay, 2 µM, is much lower than 10 µM, the concentration that is usually used for effective target knockdown for RNAi reagents. We concluded C1 or C2 forms of sliDRNA can be potent RNAi triggers and will reduce the cost by 40 to 60 percentage.

**Fig. 9:**
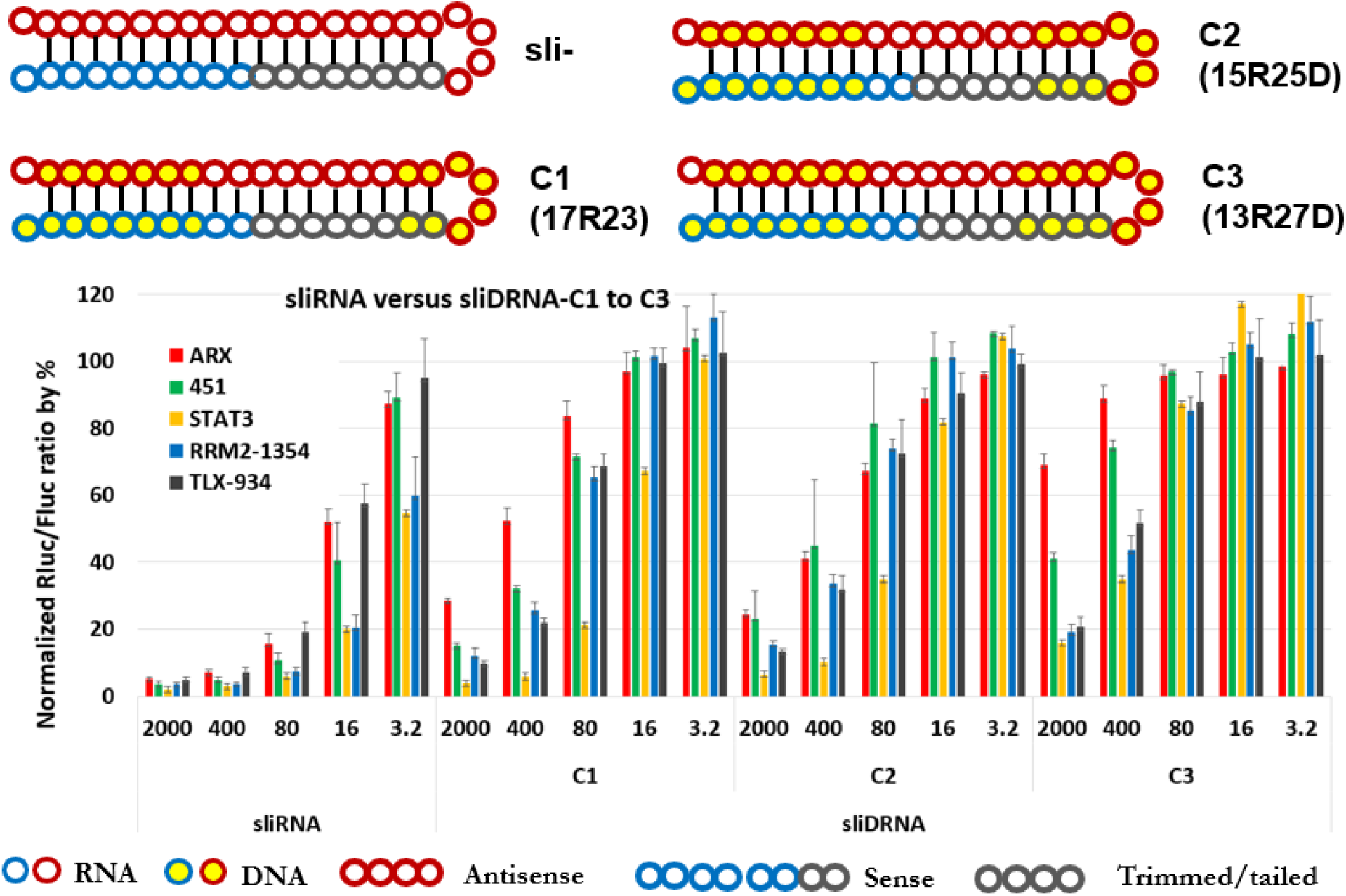
Optimal compositions of sliDRNA. a. Schematic representation of sliRNA and three provisional compositions of sliDRNA; b. Results from reporter assay.

### sliDRNA with DNA tails

Previously we have tested adding tail nts at N_40_ to check whether tailed nts have effects on sliRNA and found at least five nts can be added without significant effects^14^. Here we extend our test to extra 10 (D_41-50_) or 15 (D_41-55_) DNA nts added to the 3’ end. We tested both sliDRNA-ARX and sliRNA-451. We found there are only few percentage drops in efficacy (**Fig. 10**). We can conclude that the 3’ end of sliRNA can be used for attachments of other moieties, nts from N_31_ to N_40_ plus at least up to 15 tailed nts can be incorporated into sliRNA to create dual or multiple functional therapeutic conjugates.

**Fig. 10.**
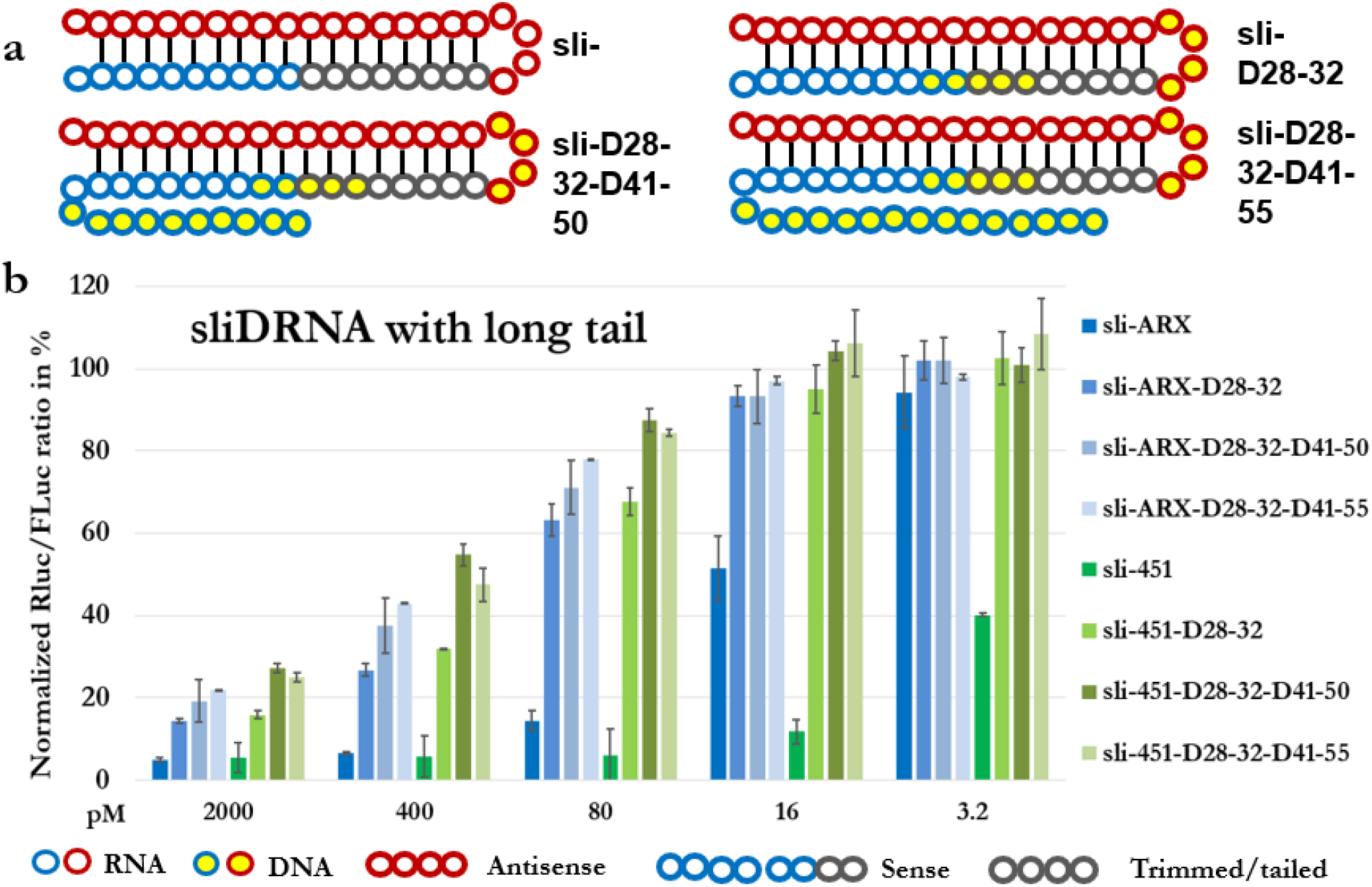
: sliDRNA with tailed nts. a. Schematic representation of all sliDRNAs; b. Results from reporter assay.

### sliDRNA depends on Ago2 for function

To further validate the sliDRNA results and rule out the silencing effects are not from other endoribonucleases, such as Ribonuclease H (RNase H), although their potency suggested that it is unlikely. We performed reporter assay in wildtype, DROSHA knockout (KO), Dicer KO and Ago2 KO HCT116 cells. While the results proved sliDRNA relies on Ago2 for function, we also observed improved silencing activities for sliDRNA in DROSHA or Dicer KO cells (**Fig. 11**). The improvement may be caused by more Ago2 are available for sliRNA under DROSHA or Dicer depletion conditions that reduced endogenous small RNA population occupy less Ago2.

**Fig. 11.**
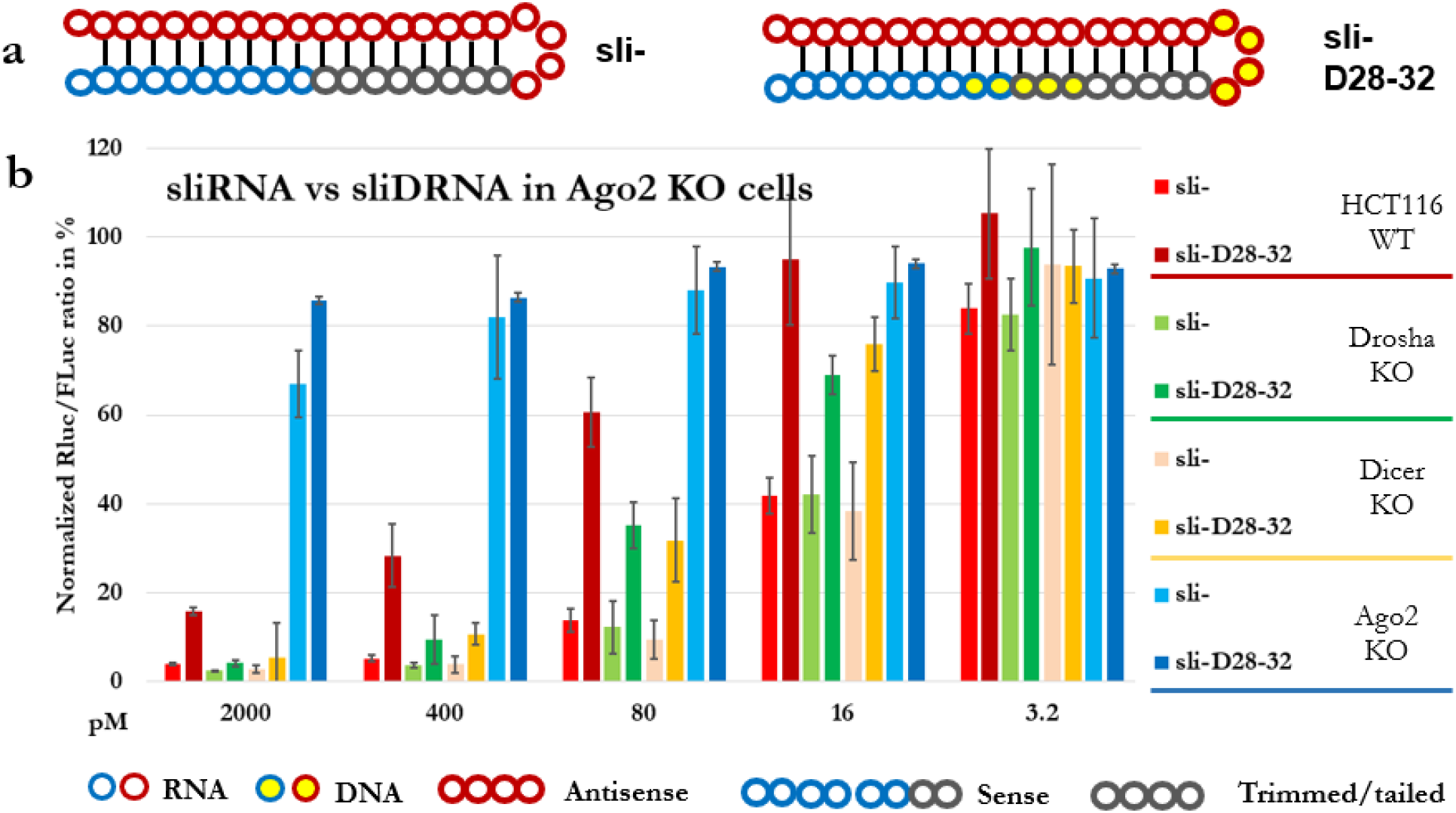
: sliDRNA function relies on Ago2. a. Schematic representation of all sliDRNAs; b. Results from reporter assay.

**Table 1:**
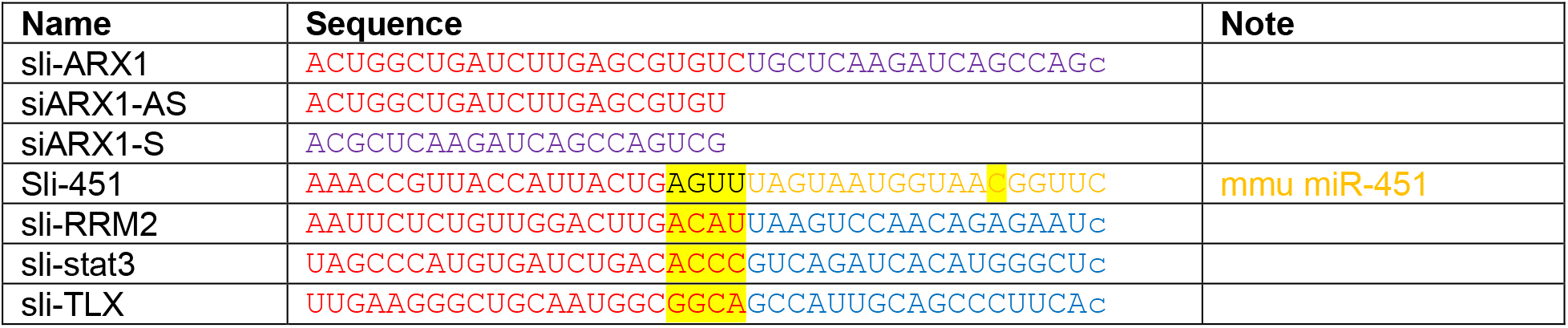
Oligo sequences used:

## Discussion and limitations

RNA interference (RNAi), mediated by small interfering RNAs (siRNAs), is a powerful tool for manipulating gene expression. Currently, there are two major types of RNAi triggers, siRNA and sliRNA, which are distinguished by their biogenesis and functional mechanisms. Previously we named Ago2 sliced RNA as agsiRNA for chemically synthetic form and agshRNA for pol III promoter expressed form^14^. From the current publication and onwards, we will named them as sliRNA and slishRNA, respectively. In the current study, we have systematically studied and characterized the nts compositions of sliDRNAs, a cost-effective step that can benefit the next stage to form the optimal composition of cm-sliRNA for sliRNA *in vivo* applications.

There is a special interest in incorporating cm-nts into traditional siRNAs due to RNA nts are significantly more unstable and prone to degradation than DNA. It is necessary to use more stable cm-nts to change their stability and to neutralize charges to enhance their transportability to facilitate *in vivo* applications. Our current studies showed DNA substitutions have different effects on siRNAs and sliRNAs due to their different activation mechanisms and sliDRNA can be potent RNAi triggers, which will save around 40% compared to sliRNA. The current study has paved the way to form an optimal composition of cm-sliRNA to expand sliRNA application soon. We showed again that the 3’ end of sliRNA has the flexibility for adding other moieties, such as extra cm-nts, RNA motifs, and delivery reagents, with minor effects on efficacy.

We extend our result derived from sli-ARX to sli-451, sli-STATs, sli-RRM2, and sli-TLX. While there are variations in silencing different genes, the sliDRNA followed the efficacy of their parental sliRNA for all three candidate sliDRNA forms, indicated the rules should be universal to design any sliDRNA.

Our study also reveals some unknown mechanisms of Ago2 slicer action. Our sliDRNA-ARX results showed sli-ARX-DR-II form, that N_13:28_ and N_14:27_ pairs are critical for sliDRNA potency. Becsaue they are A:T and G:C pairs, it is unlikely the effects are caused by special nts at these positions. Therefore, the effects on Ago2 slicer activities from N_13:28_ and N_14:27_ pairs in sliRNA deserve further study. A structure study may help reveal more insights about Ago2 slicer function, at least for sliRISC. Most likely the small hairpin structure makes Ago2 work in a different conformation in sliRISC that involved the rotation of N and PAZ domains.

We have not examined the potential effect on Ago2 cleavage rate caused by DNA nts. Previous study showed nts at N_7, 10, 17_ can affect the silencing rate for siRNA^36^. However, in sliRNA, PS also acts as a target, making it difficult to measure the rate for target cleavage. Whether nts at these positions also have the same effects for sliRNA remains a question. RISC function is also affected by other co-factors, it is also unknown whether these co-factors also perform the same function in sliRISC^37^.

smRNA-seq is the best way to characterize the processed intermediates of sliRNA after Ago2 slicing which can be used to check whether the DNA replacement affect sliDRNA processing to affect their potency or the DNA nts in the GS strand reduced the GS target binding to affect their potency. However, DNA nts substitution will affect the smRNA-seq library construction step and affect both read count and read length of processed sliDRNA, thus not reliable. We observed about 10 to 20% increased reads of miR-451 in HEK-293 cells, much less than expected spike of miR-451 reads after sliDRNA-451 transfection, despite the robust knock down rate (**Fig.10**). Therefore, the reporter assay results probably are the overall effects by DNA replacement effects on both sliDRNA processing and sliRISC function.

## Materials and Methods

### Cell lines and cell culture

HEK-293, HEK-293T, HCT-116, HCT-116 DROSHA KO, Dicer Ko, and Ago2 KO cells were maintained in high glucose (4.5 g/l) DMEM supplemented with 2 mM glutamine, 10% FBS, and 2 mM penicillin/streptomycin. Cells were incubated at 37 °C, 5% CO_2_.

### Transfection

For reporter assays, RNAi triggers and reporter constructs were co-transfected into cells by using Lipofectamine 2000 (Thermo Fisher Scientific, Grand Island, NY) as previously reported ^14^. For each experiment, at least three independent transfections were performed in duplicate in 24-well plates. Cell were grown to 75 to 85 % confluency in 500 µl medium and were transfected with luciferase reporter (50 ng), and different amounts of siRNA (100 ng of stuffer DNA, plus 1µl of siRNA stock at 1 µM, 200 nM, 40 nM, 8 nM, and 1.6 nM, and 1 µl of Lipofectamine 2000).

For qPCR analysis, the final concentration of 10 nM siRNA was transfected by RNAiMAX (Thermo Fisher Scientific, Grand Island, NY) in 12-well plates. Forty-eight hours post transfection, cells were lysed with TRIzol® (Thermo Fisher Scientific, Grand Island, NY) for total RNA isolation.

### Dual-luciferase reporter assays

To deliberate the DNA replacement studies, instead of using single dosage to measure RNAi reagents potency, we have delineated the knockdown experiments in series dilutions of RNAi reagents using reporter assay that measure the output at protein level in quantitative way. All reporter assays were performed using psiCheck 2.0-based, dual-luciferase reporter plasmids from Promega (Madison, WI) that express both *firefly* luciferase (Fluc) and *Renilla* luciferase (Rluc). Reporters carried complementary target sequences that were constructed by inserting annealed oligonucleotides into the Xho I/Spe I sites of the 3’ UTR of the Rluc gene in psiCheck2.2 vector ^14^. These reporters were used to quantify gene silencing. Forty-eight hours after transfection, cells were lysed with 100 μl passive lysis buffer (Promega) and luciferase levels for 20 μl of lysate were determined (Dual-Luciferase reporter assay kit and GloMax 96 Microplate Luminometer, Promega). Changes in expression of Rluc (target) were normalized to Fluc (internal control) and then calculated relative to the scrambled sliRNA control. The relative ratios of Rluc/Fluc were used to measure the efficiency of silencing. Data was averaged from at least three independent transfections, and each transfection had at least two replicates. Error bars indicate the standard deviation.

### qPCR

BIO-RAD (Hercules, California) iTaq™ Universal SYBR® Green One-Step Kit was used for qPCR. Briefly, total RNA was isolated by TRIzol followed by DNase treatment. In each reaction, 500 ng DNase treated total RNA was used, and all other reagents were used as specified in the protocol provided in the kit. We also followed the qPCR program suggested by the vendor. GAPDH gene was used as a normalization control to calculate ΔCt for each sample. 2^−ΔΔCt^ value of each sample (ΔΔCt = ΔCt_siRNA_-ΔCt_siRNA ctrl_) was used for RNAi knockdown measurement and converted to %. 2^−ΔΔCt^ value of each sample (ΔΔCt = ΔCt_siRNA treated HEK293 cells_ –ΔCt_untreated HEK293 cells_) was used for calculating fold changes in gene expression to measure immune response. Data were averaged from at least three replicates. Error bars indicate the standard deviation of the mean.

### Oligonucleotides

All oligonucleotides were synthesized by Integrated DNA Technologies (Coralville, Iowa).

## Acknowledgments

Authors would like to thank Dr. Marcus Peter from Northwestern University for generously proving us DROSHA and Dicer KO HCT 116 cells.

## Author Contributions

GS and AR conceived and designed experiments. GS, and EL performed experiments. GS and YS drafted the final version of the manuscript.

